# Time is vision: functional preservation and enhanced capacity for recovery in subacute occipital stroke

**DOI:** 10.1101/847954

**Authors:** Elizabeth L. Saionz, Duje Tadin, Michael D. Melnick, Krystel R. Huxlin

## Abstract

Stroke damage to the primary visual cortex (V1) causes a loss of vision known as hemianopia or cortically-induced blindness (CB). While early, spontaneous, perimetric improvements can occur, by 6 months post-stroke, the deficit is considered chronic and permanent. Despite evidence from sensorimotor stroke showing that early injury responses heighten neuroplastic potential, to date, rehabilitation research has focused only on chronic CB patients. Consequently, little is known about the functional properties of subacute, post-stroke visual systems, and whether they can be harnessed to enhance visual recovery. Here, for the first time, we show that *conscious visual discrimination* abilities are partially preserved inside subacute, perimetrically-defined blind fields, disappearing by 6 months post-stroke. Complementing this discovery, we show that global motion discrimination training initiated subacutely leads to comparable magnitude of recovery as that initiated in chronic CB. However, it does so 6 times faster, generalizes to deeper, untrained regions of the blind field, and to other [untrained] aspects of motion perception, preventing their degradation upon reaching the chronic period. Untrained subacutes exhibited only spontaneous improvements in perimetry - spontaneous recovery of motion discriminations was never observed. Thus, in CB, the early post-stroke period appears characterized by gradual - rather than sudden - loss of visual processing. Subacute training stops this degradation, and is dramatically more efficient at eliciting recovery than identical training in the chronic period. Finally, spontaneous improvements in subacutes appear restricted to luminance detection, whereas recovering discrimination abilities requires deliberate training. Simply stated, after an occipital stroke, “*time is VISION*”.

**One Sentence Summary:** The first 3 months after an occipital stroke are characterized by a gradual - not sudden - loss of visual perceptual abilities and increased rehabilitative potential if visual discrimination training is administered in the blind field.

## Introduction

The saying *“time is brain”* after stroke may be true, but if that stroke affects primary visual cortex (V1), the urgency seems to be lost. Such strokes cause a dramatic, contralesional loss of vision known as hemianopia or cortically-induced blindness (CB) (*1, 2*). Our current understanding of visual plasticity after occipital strokes or other damage is largely informed by natural history studies that show limited spontaneous recovery early on, with stabilization of deficits by six months post-stroke (*2–5*). By that time, CB patients are considered “chronic” and exhibit profound visual field defects in both detection and discrimination contralateral to the V1 damage (*6–10*). In fact, research into post-stroke visual function and plasticity has focused on patients in this chronic phase (*11*), precisely because of the stability of their visual field defects. However, therapeutically, this approach runs counter to practice in sensorimotor stroke, where it has been shown that early injury responses heighten neuroplastic potential (*12–19*).

Although some have argued that the visual system is not capable of functional recovery in the chronic phase post-stroke (*20–22*), multiple studies, from several groups worldwide, have shown that gaze-contingent visual training with stimuli presented repetitively inside the perimetrically-defined blind field can lead to localized visual recovery – both on the trained tasks and on visual perimetry measured using clinical tests (*7, 9–11, 23–26*). However, the training required to attain such recovery is intense and lengthy (*9–11*), and recovered vision appears to be low-contrast and coarser than normal (*9, 10, 26*). It is currently unknown why visual recovery in chronic CB is so difficult, partial, and spatially restricted. Possible explanations include that V1 damage kills a large portion of cells selective for basic visual attributes such as orientation and direction, and that it causes a shift in the excitation/inhibition balance in residual visual circuitry towards excessive inhibition (*27*). Excessive intra-cortical inhibition can limit plasticity and raise the threshold for activation of relevant circuits. These factors may explain why training that starts more than six months post-stroke is arduous and why recovery is incomplete.

Thus, while there is much left to do to improve rehabilitation strategies for the increasingly large population of chronic CB sufferers (*1*), the situation is much worse for acute and subacute CB. Indeed, this group of early post-stroke patients has been almost completely ignored in the vision science literature. There are no published, detailed assessments of visual function within CB fields in the first few weeks after stroke, nor do we know anything about the potential for training-induced recovery during this early post-stroke period. This is in marked contrast to sensorimotor stroke, which has been investigated as soon as five days after ischemic damage, with current treatment guidelines advocating early rehabilitation to facilitate greater, faster recovery (*12–15*). In addition to the resolution of stroke-associated inflammation and swelling, this early period is characterized by a shift in the excitation/inhibition balance towards excitation, up-regulation of growth and injury response factors (especially brain-derived neurotrophic factor), changes in neurotransmitter modulation (especially GABA, glutamate and acetylcholine), and even re-emergence of a critical-period-like state (*16–19*). These cellular and environmental cerebral changes could in fact underlie the observed spontaneous recovery in luminance detection perimetry in the first few months after visual cortical strokes (*2, 6, 28, 29*). Moreover, structural barriers to plasticity have yet to form, in particular myelin-related proteins inhibiting axonal sprouting, perineuronal nets of chondroitin sulfate proteoglycans, and fibrotic tissue (*16, 17*). Thus, whether this post-stroke *critical period* could be recruited to attain greater and faster recovery of visual functions in the blind field is the second question asked here.

## Results

We began by measuring the basic properties of vision in 18 subacute CB fields (mean±SD=7.7±3.4 weeks post-stroke, range 2.0-12.7 weeks), comparing them with performance in the intact visual field of the same participants, and in 14 adults with chronic CB (33.4±54.5 months post-stroke, range 5-226 months). Subject demographics are detailed in **Table 1**. Brain scans illustrating individual lesions and visual field defects are presented in **Figs 1** and **2**.

**Fig. 1.**
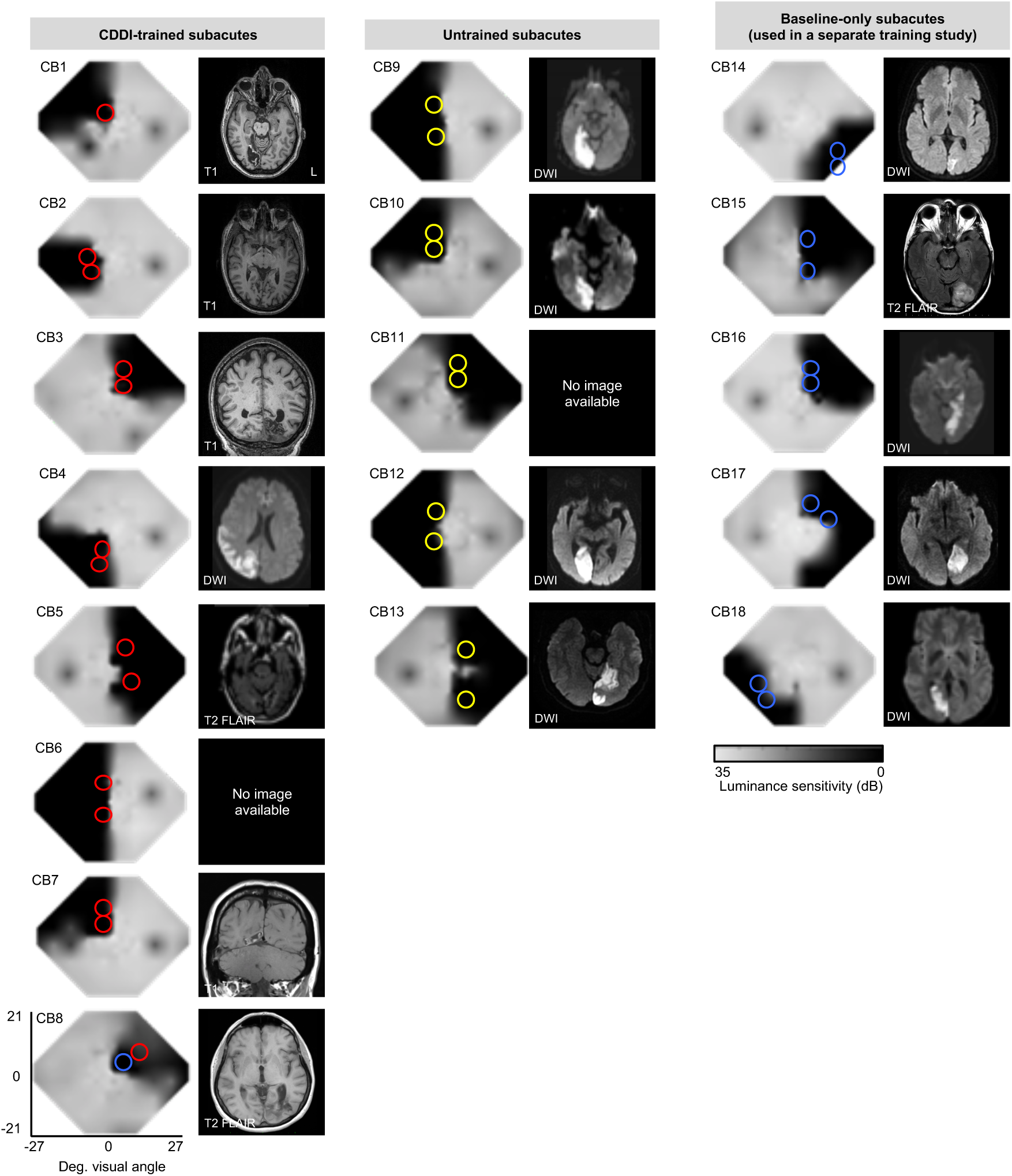
Baseline Humphrey visual field composite maps, magnetic resonance images (MRI) and testing/training locations in subacute participants. Grey scale denoting Humphrey-derived visual sensitivity is provided under right-most column. MRI type (T1, diffusion-weighted imaging [DWI], T2-weighted fluid-attenuated inversion recovery [T2-FLAIR]) is indicated on radiographic images, which are shown according to radiographic convention (left brain hemisphere on image right). Red circles: CDDI training locations; yellow circles: putative training locations in untrained controls, which were only pre- and post-tested; blue circles: locations tested at baseline in subacutes who were used in a separate training study (designated “other” training type in **Table 1**).

**Fig. 2.**
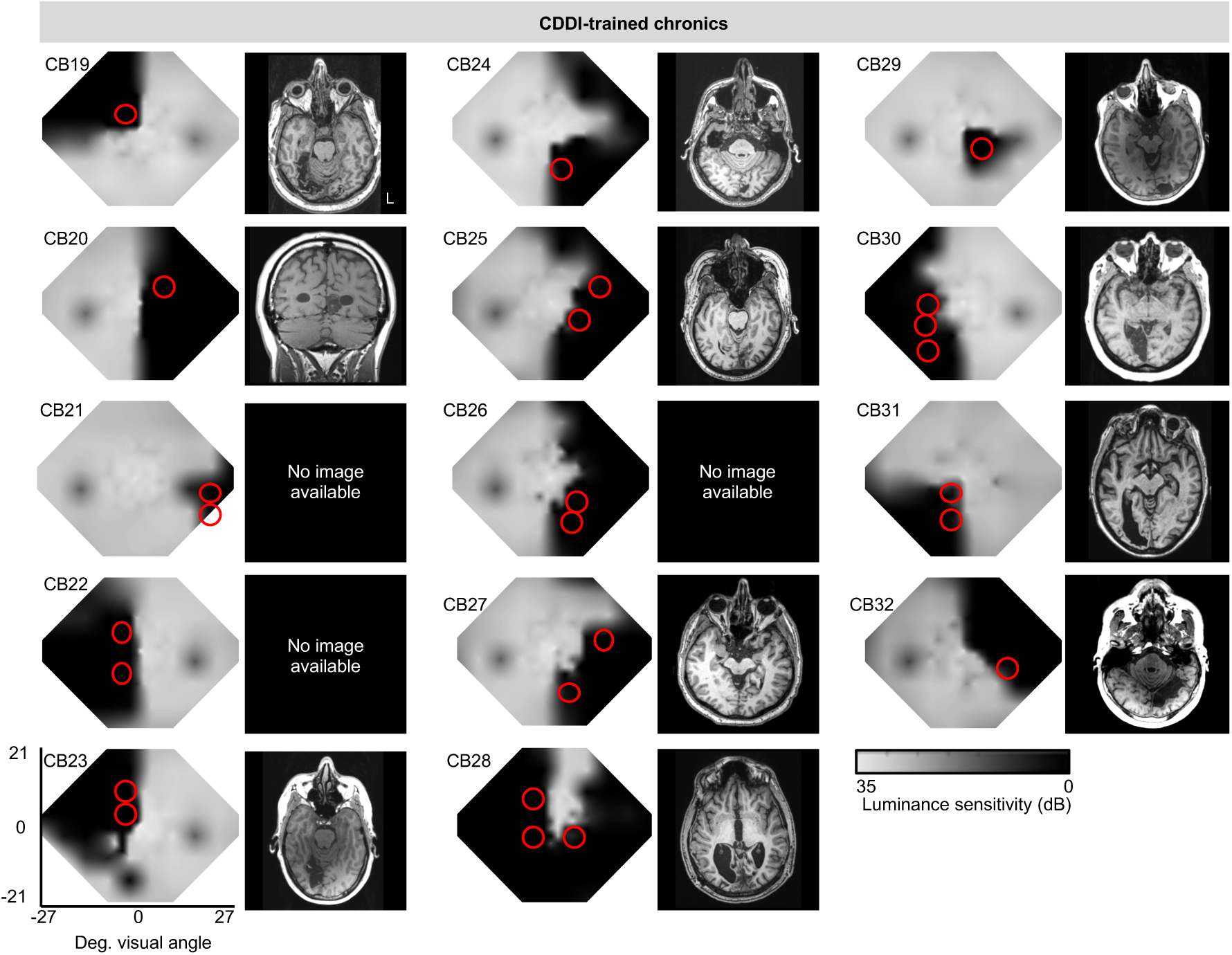
Baseline Humphrey visual field composite maps, structural (T1) magnetic resonance images and training locations in chronic participants. Grey scale denoting Humphrey-derived visual sensitivity is provided under right-most column. MRIs are shown according to radiographic convention with left brain hemisphere on the right-hand side of the image (L). Red circles: CDDI training locations. Data from these chronic subjects were previously published (*7*).

**Table 1.**
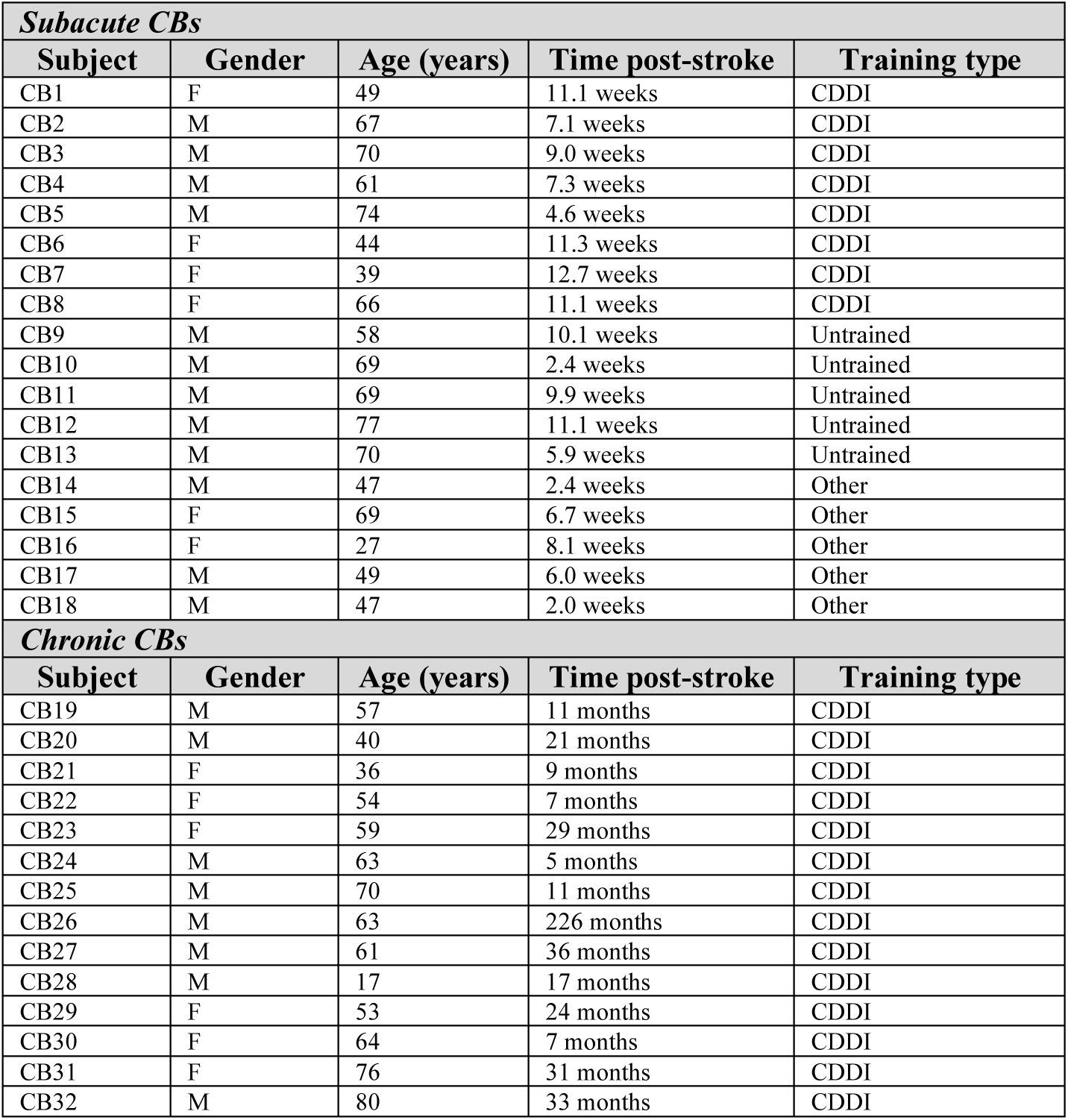
Participant demographics, testing and training parameters. M, male; F, female; CDDI, coarse direction discrimination and integration training. Patients CB14-18 underwent “other” types of training – their outcomes are part of a different study.

Visual field deficits were first estimated from Humphrey visual perimetry, as previously described (*7*) and served as a starting point to precisely map the position of the blind field border and select training locations. This mapping was done with a coarse (left/right) global direction discrimination and integration (CDDI) task and a fine direction discrimination (FDD) task, both using random dot stimuli (*7, 9, 10, 26*). Suitable, initial training locations were picked from these test results according to previously-published criteria (*9, 10, 26*) and are shown to scale, with colored circles superimposed on composite Humphrey visual maps in **Figs. 1** and **2**.

At each selected, training location in the blind field, luminance contrast sensitivity functions (CSFs) were then estimated for both static, non-flickering, orientation discrimination and direction discrimination. Finally, intact field performance was also collected in each participant: for each task, performance was measured at locations mirror-symmetric to the blind field locations selected for initial training. This was essential to provide a patient-specific, internal control for “normal” performance on each task.

### Preserved motion discrimination in subacute but not chronic blind fields

Consistent with prior observations, within their perimetrically-defined blind fields, our chronic CB participants failed to discriminate opposite motion directions (*10, 26, 30*) or to effectively integrate across motion directions (*7, 9, 10*) (**Fig. 3a-b**) – tasks that elicited normal, threshold levels of performance at corresponding locations within their intact visual hemifields. Subjects verbally-reported detecting that visual stimuli had been briefly presented within their blind fields, but they could not discriminate the global direction information contained in these stimuli above chance-level performance (50% correct in these two-alternative, forced choice tasks).

**Fig. 3.**
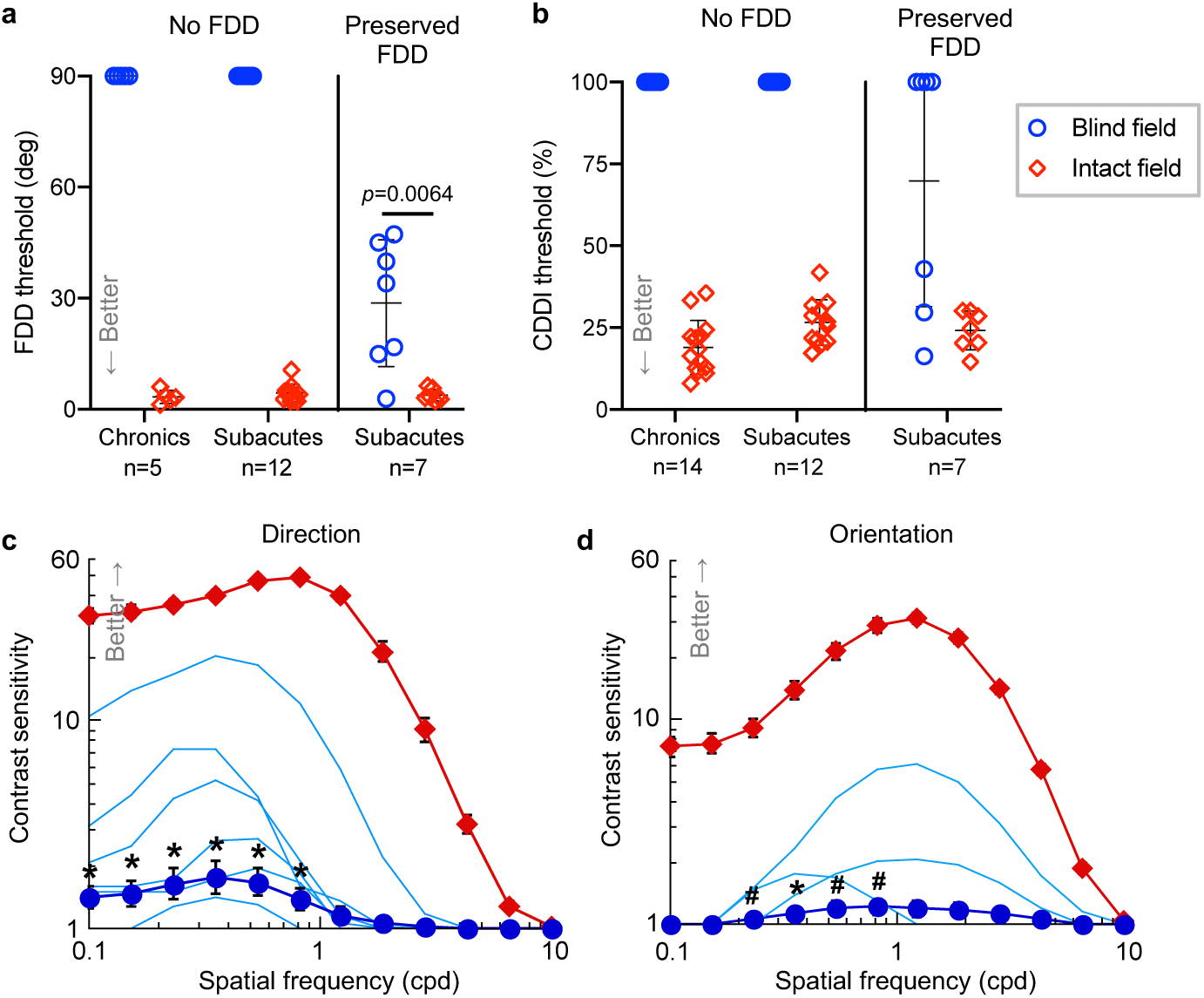
Preserved visual discrimination abilities in subacute but not chronic CB fields. **a**, Plot of individual baseline FDD at blind-field training locations and corresponding, intact-field locations in chronic and subacute CBs. Bars indicate means±SD. Baseline FDD was unmeasurable in all chronics and 2/3 of subacutes, but measurable in 1/3 of subacutes. CB15 is included in both subacute categories because of ability to discriminate FDD in one blind-field quadrant but not the other, illustrating heterogeneity of perception across CB fields. As a group, subacutes’ baseline FDD thresholds were better than chronics’ (1-sample t-test versus mean of 90°, t_18_=3.08, p=0.0064). However, subacutes with preserved FDD had worse thresholds than in their own intact hemifields (paired t-test, t_6_=4.09, p=0.0064). **b**, Plot of baseline CDDI at blind- and corresponding intact-field locations in chronic and subacute CBs, stratified by preservation of blind-field FDD. Three subacutes with preserved blind-field FDD had measurable CDDI thresholds, a phenomenon never observed in chronics. **c**, Baseline CSFs for direction discrimination in the blind and intact fields of subacutes (data points=means±SEM); light blue lines denote CSFs of participants with preserved blind-field sensitivity (significant in n=5, p<0.005; in the 6^th^ subject, p=0.16, see Methods for bootstrap analysis). Group t-tests were performed at each spatial frequency: * p<0.05, # p<0.10. There were significant effects for peak CS (t_14_=2.38, p=0.016) and total area under the CSF (t_14_=2.10, p=0.027). **d**, Baseline CSFs for orientation discrimination in subacutes. Labeling conventions as in (c). Statistics for peak CS and area under the CSF: t_13_=1.5, p=0.079 and t_13_=1.62, p=0.065, respectively.

In contrast, when performing the same tasks, just over a third of subacutes could still perceive and discriminate relatively fine direction differences (FDD) at multiple locations in their blind fields, albeit with FDD thresholds usually poorer than at equivalent locations in their own intact fields (29±17° *versus* 4±2°) (**Fig. 3a**). A further 43% of these subacutes could also integrate motion direction in their blind field, with CDDI thresholds approaching their own intact-field levels (**Fig. 3b**). When performance exceeded chance, participants always reported subjective awareness of the stimulus and a clear sensation of motion (in a direction above or below the horizontal meridian for the FDD task, and left-or rightward for the CDDI task). In contrast, subacutes who performed at chance on the FDD task in their blind field also performed at chance when asked to integrate motion direction into a discriminable percept (CDDI task). These participants could usually detect appearance and disappearance of the visual stimuli, but like chronic patients, were unable to reliably identify the global direction of motion they contained.

Even more surprising than the preservation of global motion discrimination measured using high-contrast, random dot stimuli, a third of subacutes tested also had preserved contrast sensitivity for opposite direction discrimination of small Gabor patches in their blind field (**Fig. 3c**). At the same blind-field locations, preserved contrast sensitivity for the orientation discrimination of static, non-flickering Gabors was only observed in 3 of these participants (**Fig. 3d**). As with the random dot stimuli, where participants had preserved contrast sensitivity, they reported sensation of the stimulus and its direction/orientation. To our knowledge, even partial preservation of luminance contrast sensitivity in perimetrically-defined blind fields has *never* been described in the literature on this patient population. That contrast sensitivity should be preserved at all is also somewhat surprising, since it was measured *within* perimetrically-defined blind fields, and Humphrey perimetry is in essence, a measure of luminance contrast detection. However, there are some key differences between Humphrey stimuli and those used to measure contrast sensitivity in the present experiments: Humphrey uses much smaller spots of light than the 5°-diameter Gabor patches in our psychophysical tests, and it uses luminance increments in these small spots relative to a bright background. The larger Gabors may have been more detectable by invoking spatial summation (*31–33*), which would have improved perceptual performance.

Nonetheless, that some subacute CBs should possess measurable contrast sensitivity functions in their blind fields is in stark contrast to the lack of such functions in chronic CBs. Over more than a decade of testing, we and others have consistently found luminance contrast sensitivity – prior to training interventions - to be un-measurable in such patients (*8–10*).

In sum, we report here the discovery of preserved direction discrimination, direction integration abilities and even luminance contrast sensitivity [strongest for direction discrimination] in the perimetrically-defined blind fields of a significant proportion of subacute participants less than 3 months post-stroke. Given that threshold-level performance is *never seen* in chronic CB participants, we posit that subacutes retain functionality of key visual circuits, which are then lost by the chronic period. It also highlights the highly modular nature of visual processing, which allows even extensive brain damage to occur without eradicating all visual function, at least initially.

### Effect of CDDI training

We next turned our attention to the as-yet-unaddressed question of how subacute CB patients respond to visual training. To ensure a fairer comparison with chronic CBs, who have no preserved discrimination abilities in their blind field at baseline (*9, 10, 26*), we sub-selected those subacutes with deficits in global motion perception (CB1-13) in their blind field (**Fig. 1**). These subacutes were alternatingly assigned, in the order they were enrolled, to either CDDI training or no training until 5 people were enrolled in each. All subsequently enrolled subacutes (n=3) were directed into the training group, so that we ended up with a total of 5 untrained (yellow circles in **Fig. 1**) and 8 trained CB patients (red circles in **Fig. 1**). Training was performed until subacutes entered the chronic (>6 months) post-stroke period, and their data were compared with previously-published data (*7*) from 14 CDDI-trained, chronic CB participants (**Table 1, Fig. 2**). Daily, in-home training on the CDDI task was administered over a comparable number of sessions, each 300 trials long (subacutes: 125±46 sessions, chronics: 129±82 sessions). Training protocols were identical to those used previously in chronics, with initial training locations selected where participants could not integrate motion direction sufficiently well to attain a measurable CDDI threshold (*7, 9, 10*). Trained subacute participants returned for in-lab testing using eye-tracker-enforced, gaze-contingent stimulus presentations just after they entered the chronic stage (7.0±0.9 months post-stroke, after 4.8±1.1 months training). Untrained subacutes were brought back when they reached 7.9±3.7 months post-stroke.

Training improved CDDI thresholds comparably across all trained participants, both subacute and chronic (**Fig. 4a, b**), but surprisingly, untrained subacutes exhibited *no spontaneous recovery of CDDI thresholds in their blind field* - they remained unable to integrate motion direction at all pre-tested, blind-field locations (**Fig. 4b**).

**Fig. 4.**
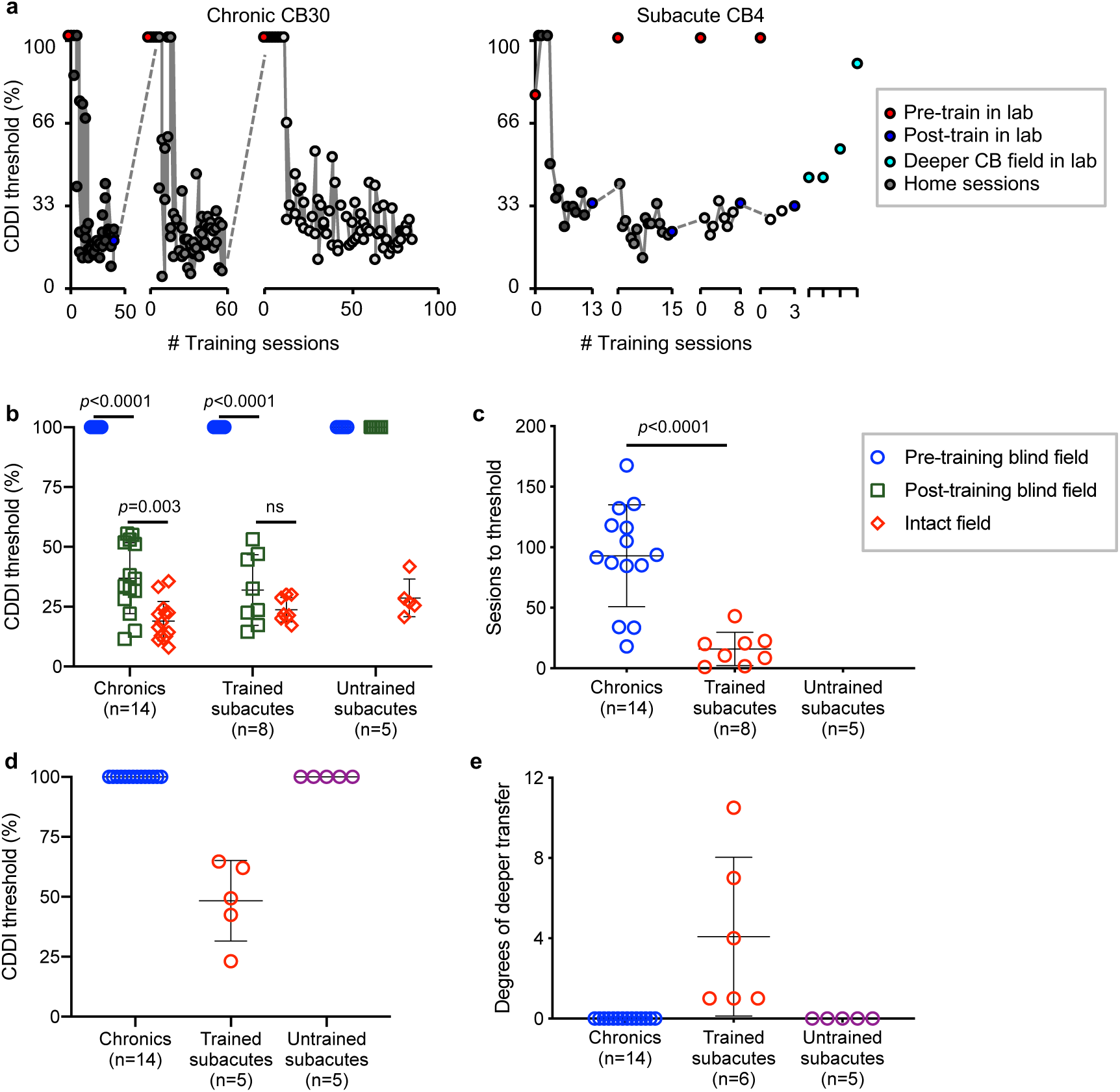
Trained subacutes recover direction integration faster and deeper than chronics. **a**, Training data for representative chronic and subacute CBs. **b**, Plot of individual CDDI thresholds at training locations pre- and post-testing. Bars indicate mean±SD. Two-way repeated measures ANOVA for group (chronics, trained subacutes, untrained subacutes) across locations (pre-training blind field, post-training blind field, intact field) was significant: F_4,48_=37.11, p<0.0001. Post-hoc Tukey’s multiple comparisons tests within group are shown on graph. **c**, Plot of the number of training sessions to reach normal CDDI thresholds in the blind field. Bars indicate mean±SD. Chronics required significantly more training sessions than trained subacutes: unpaired t-test t_20_=4.98, p<0.0001. **d**, Plot of initial CDDI threshold at location 1° deeper into the blind field than trained/tested location. Bars indicate mean±SD. Only trained subacutes had measurable thresholds deeper than the trained blind-field location. **e**, Plot of degrees of visual angle by which random dot stimulus could be moved deeper into the blind field than the last training/testing location, while still able to attain a measurable CDDI thresholds. All trained subacutes had measurable CDDI thresholds deeper into the blind field – something never observed in chronic or untrained subacutes. Bars indicate mean±SD. Two trained subacutes were not included due to extent of recovery exceeding our ability to measure depth in the blind field. One-way ANOVA across groups F_2,22_=10.69, p<0.0001.

Among trained participants, a key difference also emerged: subacutes recovered significantly faster than chronics, reaching normal, stable CDDI thresholds in only 16±14 training sessions per blind-field location, compared to 93±42 sessions in chronics (**Fig. 4c**). Additionally, subacute CBs exhibited generalization of CDDI recovery to untrained blind-field locations (**Fig. 4a**, right panel; **Fig. 4d,e**), something *never* seen in chronic CBs (**Fig. 4a**, left panel; **Fig. 4d,e**) (*9, 10*). By the end of training, subacutes and chronics had regained global motion discrimination on average 4° and 0° deeper into the blind field than the deepest trained location, respectively (**Fig. 4e**). Untrained subacutes, as expected, showed no significant motion perception improvements either at the originally-tested locations or deeper into their blind field.

### Generalization to untrained tasks

#### Fine direction discrimination

subacutes selected for CDDI training fell into 2 categories: those with and without preserved FDD thresholds. As such, we considered outcomes in these two sub-groups separately. Among subacutes with no baseline preservation of FDD in the blind field, CDDI training transferred to and improved FDD thresholds in 4/6 participants (**Fig. 5a**). All trained chronics (who never have preservation of FDD at baseline) also exhibited transfer of learning to FDD but they attained better FDD thresholds than subacutes. Given that longer training tends to enhance learning transfer (*34*), this better outcome in chronics could be related to the subacutes spending less time training at each blind-field location because of their faster learning rate. Nonetheless, consistent with our prior studies (*10, 26, 30*), neither subacutes nor chronics achieved intact-field levels of FDD thresholds following CDDI training. Notably, untrained subacutes exhibited no spontaneous recovery of FDD thresholds (**Fig. 5a**). Finally, in subacutes with preserved blind field FDD thresholds at baseline, CDDI training maintained but did not further enhance FDD thresholds (**Fig. 5b**). As such, blind field CDDI training in the subacute period may have been critical for preserving existing FDD performance. This was further evidenced in untrained subacute CB10, who could discriminate fine direction differences in the blind field at baseline; however, by 6 months post-stroke, absent any training, this ability was lost and CB10 performed like any typical, untrained, chronic CB patient (*10, 26*).

**Fig. 5.**
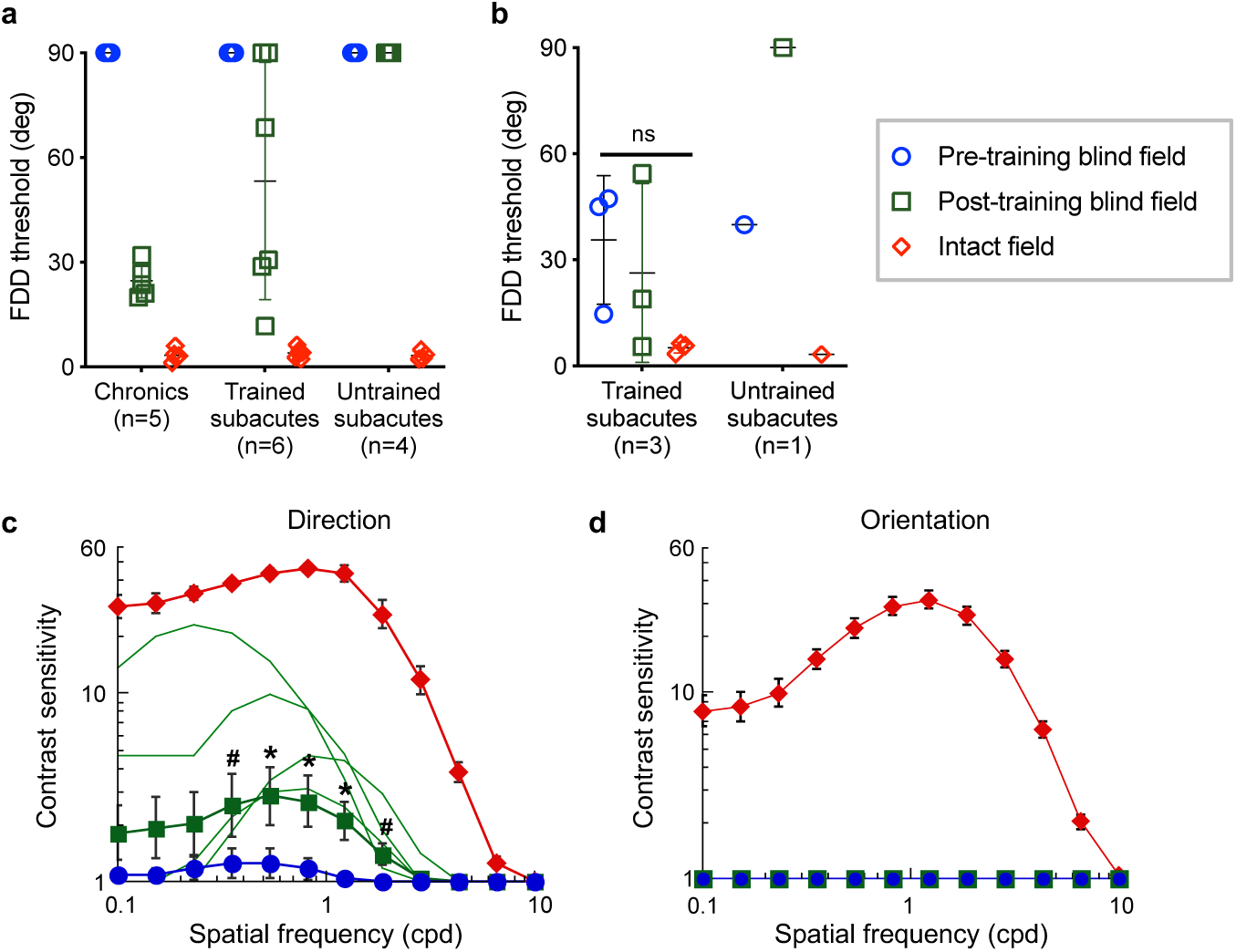
Subacute training on CDDI improves FDD thresholds and motion CSFs at trained, blind-field locations. **a**, Plot of FDD thresholds in participants without baseline FDD, before and after CDDI training. Labeling conventions as in **Fig. 4b**. CDDI training improved FDD thresholds in most cases, whereas untrained subacutes never improved. A 3_participant type_x3_visual field location_ repeated measures ANOVA showed a main effect of participant (F_2,12_=9.715, p=0.0031), visual field location (F_1.007,12.09_=168.6, p<0.0001, Geisser-Greenhouse epsilon=0.5036), and a significant interaction between the two (F_4,24_=9.629, p<0.0001). Mean recovered FDD thresholds were better in chronic than subacute trained participants (Tukey’s multiple comparisons test: p<0.01). **b**, Plot of FDD thresholds in participants with preserved baseline FDD, before and after CDDI training. No enhancements in FDD thresholds were noted (1-way repeated measures ANOVA: F_2,8_=3.81, p=0.12). When left untrained, FDD thresholds worsened to chance in the 1 subacute in this group. **c**, Post-training CSFs for direction in the blind and intact fields of trained subacutes. Labeling conventions as in **Fig. 3c** except for light green lines denoting individual, post-training CSFs. CDDI training improved CS for motion direction across multiple spatial frequencies in 4/7 subacutes (n=4, p<0.010, see Methods for bootstrap analysis). Group t-tests were performed at each spatial frequency, with *p<0.05, #p<0.10. There were significant effects for peak CS (t_6_=2.45, p=0.025) and area under the CSF (t_6_=2.28, p=0.032). **d**, Post-training CSFs for orientation in subacute participants showing no improvements after CDDI training (p>0.2). Labeling conventions as in **(c)**.

#### Luminance contrast sensitivity

CDDI training in the subacute period improved luminance contrast sensitivity for direction, but not for orientation discrimination of static [non-flickering] Gabors.

Among subacutes selected for CDDI training, none had measurable baseline CS for static orientation discrimination, and only one had measurable baseline CS for direction discrimination. After CDDI training, 4/7 of tested participants achieved measurable motion CSFs (**Fig. 5c**), but orientation CSFs remained flat (**Fig. 5d**). Untrained subacutes failed to improve on either measure of CS, which remained flat. Improvement on motion CS in this training study was thus completely dependent on CDDI training.

#### Humphrey perimetry

an unexpected finding in the present study was that CDDI-trained and untrained subacutes exhibited similar changes in Humphrey perimetry (**Fig. 6a, b**). No significant differences were observed between these two groups using four separate metrics: (1) perimetric mean deviation (PMD – **Fig. 6c**), (2) area of the deficit encompassed by the 24-2 Humphrey (**Fig. 6d**), (3) area of the 24-2 Humphrey that improved by more than 6dB (**Fig. 6e**), and (4) area of the 24-2 Humphrey that lost more than 6 dB of sensitivity (**Fig. 6f**). The lack of significant improvement in Humphrey perimetry in trained subacutes is consistent with their lack of improvement in blind-field static contrast sensitivity, and illustrates a clear dissociation between training-dependent recovery of motion discriminations and spontaneous, training-independent improvements in luminance detection perimetry.

**Fig. 6.**
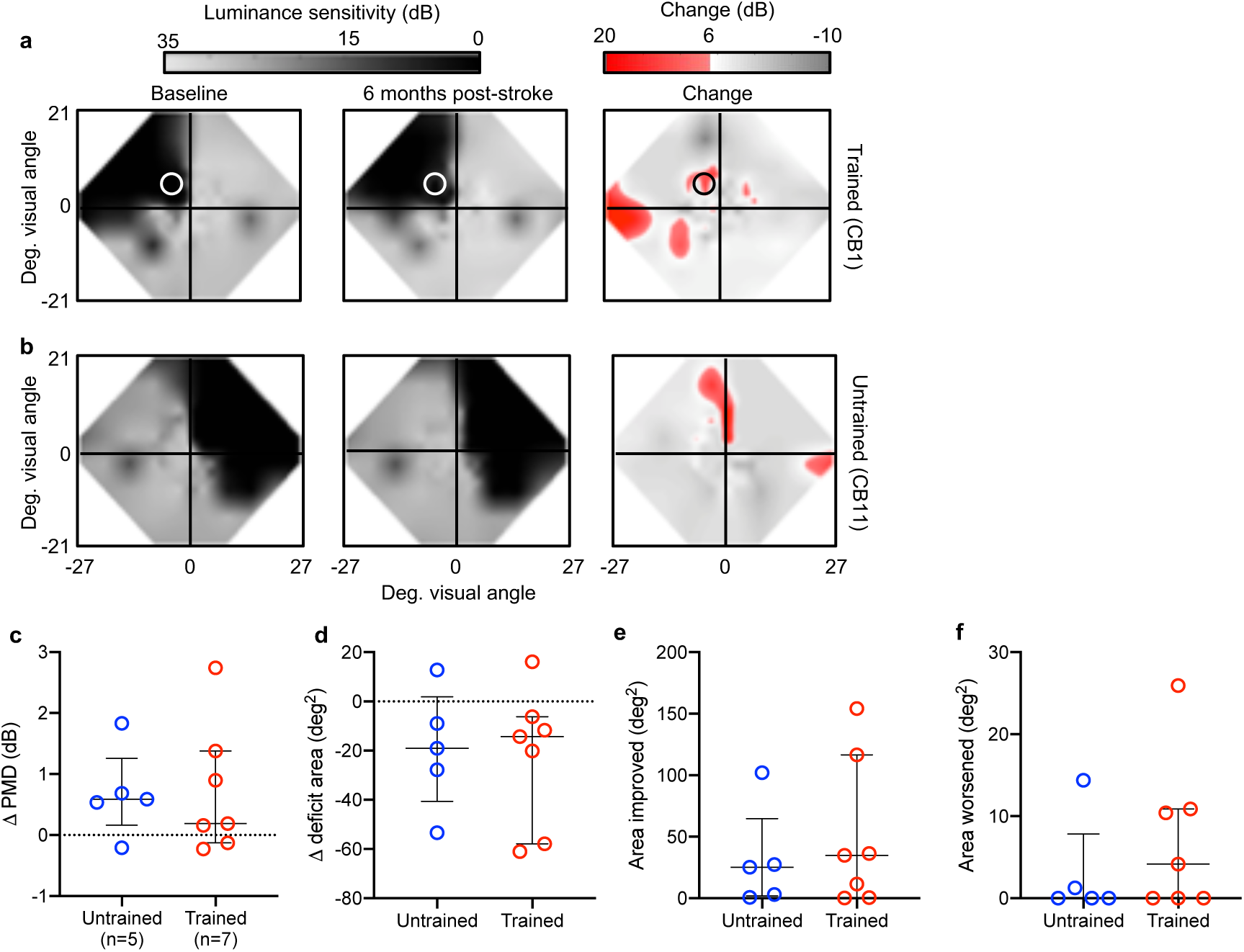
Subacute CDDI training does not improve Humphrey perimetry above spontaneous levels of recovery. **a**, Composite visual field maps of representative trained subacute participant (CB1) at baseline and post-training, along with a map of the net change in visual sensitivity (red shading), with a threshold for change of 6 dB. Training location indicated by a white circle. **b**, Composite visual field maps of representative untrained subacute participant (CB11) at baseline and follow-up, along with a map of net change in visual sensitivity. **c**, Plot of changes in the Humphrey-derived perimetric mean deviation (PMD) of individual subacute CBs who were untrained *versus* CDDI-trained. The PMD is the overall difference in sensitivity between the tested and expected hill of vision for an age-corrected, normal population. Bars indicate means±SD. No significant differences were observed between groups (independent Student’s t-test: p>0.05). **d**, Plot of change in visual deficit area in the same participants as in **c**, computed from Humphrey perimetry as previously described (*7*). No significant differences were observed between trained and untrained subacutes (independent Student’s t-test: p>0.05). **e**, Plot of the area of the Humphrey visual field that improves by >6 dB (*7*) in the same participants as in **c, d**. No significant differences were observed between trained and untrained subacutes (independent Student’s t-test: p>0.05). **f**, Plot of the area of the Humphrey visual field that worsens by >6 dB (*7*) in the same participants as in **c, d** and **e**. No significant differences were observed between trained and untrained subacutes (independent Student’s t-test: p>0.05).

## Discussion

The present study represents the first systematic assessment of visual discrimination abilities within perimetrically-defined, cortically-blinded fields in 18 subacute, occipital stroke patients. We found that – unlike chronic CBs – subacute CBs often retain global direction discrimination abilities, as well as luminance contrast sensitivity for direction. Moreover, the residual, conscious visual processing in perimetrically-blind, subacute visual fields disappears by the chronic period. That the preserved vision is consciously accessible to the patients was evident both because of their verbal reports, accurately describing the visual stimuli presented in their blind fields, and because their performance on the two-alternative forced-choice tasks used here far exceeded chance levels, going so far as to generate reliable, measurable discrimination thresholds (*35, 36*), despite a relatively short stimulus presentation (*37*) and in some cases, varying stimulus contrast (*38*).

### Substrates of preserved vision and mechanisms of vision restoration

In chronic CB, it has been hypothesized that training-induced restoration of visual motion capacities could be mediated by “alternative” visual pathways, such as those claimed to mediate “blindsight”, that proceed from the retina to the dorsal lateral geniculate nucleus (dLGN) of the thalamus, the superior colliculus, pulvinar and thence to MT and other extrastriate areas (*39–42*). With respect to the dLGN, these pathways are thought to involve primarily koniocellular projections, rather than parvocellular or magnocellular ones, because of the characteristic spatial frequency preferences exhibited in blindsight (*41, 43*). Moreover, koniocellular dLGN neurons may be less likely to undergo trans-synaptic retrograde degeneration after V1 stroke (*44, 45*), and have thus been viewed as viable conduits for visual information in longstanding CB.

Alternatively, the preserved vision in subacute CB could be mediated by spared regions of cortex in V1 (*46–50*), which may become quiescent without targeted use by the chronic period. The progressive silencing of these networks may be the result of trans-synaptic retrograde degeneration coupled with a sort of “visual disuse atrophy”, whereby weak, surviving connections getting pruned and/or down-weighted over time, as patients learn to ignore the less salient visual information within corresponding regions of their blind fields. That this vision is mediated by V1 itself is supported by the relative preservation of contrast sensitivity within the perimetric blind field, with the CSFs more reflective of the contrast response of V1 (*51–53*) than MT, which saturates at low levels of contrast (*54, 55*) and shifts to higher spatial frequencies as contrast increases (*56*). After stroke, without intervention, over a matter of months the substrates of this initially-preserved vision appear to be lost. Chronic CBs may thus differ from subacutes not only in the permissiveness of the environment around their lesion for plasticity (*16–19*), but also in the availability of neuronal substrates to perform targeted visual discriminations.

While the natural course of CB is for the residual vision of subacutes to disappear by the chronic phase after stroke, training appears to both prevent loss of remaining visual abilities and strengthen them. Here, too, the progress of vision training in subacutes (*versus* chronics) points towards greater involvement of residual V1 circuits. Performance in subacute CB fields more closely resembles that in V1-intact controls both in terms of the faster timescale of perceptual learning on the CDDI task (*57*), and the ability to transfer learning/recovery to untrained tasks, including to motion contrast sensitivity (*55, 58*). Additionally, CDDI training also improved fine direction discrimination in subacutes around a motion axis orthogonal to that trained (since the FDD task involved global motion discrimination along the vertical axis). Such transfer (both to untrained directional axes and FDD thresholds) was previously reported for chronic CB patients who trained with CDDI (*10, 26*). However, the subacutes trained on CDDI in the present study did not transfer to FDD as consistently as chronics. As mentioned earlier, generalizability of learning may have been sub-optimal in the presently-tested subacutes because of their faster learning rates and the resulting shorter time spent training per blind-field location (*34*).

### Dissociation between visual detection (perimetry) and discriminations

An important dissociation in the present results pertained to the visual behavior of untrained subacutes. As predicted, this group of 5 participants sustained measurable, spontaneous improvements in their Humphrey visual fields (∼0.5 dB increase in PMD, over ∼25 deg^2^), largely located around the blind fields’ borders, as previously reported for chronic patients (*7*). However, the same, untrained subacutes did not recover visual discrimination abilities (CDDI, FDD or contrast sensitivity) at any of the pre-tested blind-field locations, even if these locations were within perimetrically-defined border regions that exhibited spontaneously-improved Humphrey sensitivity (e.g. see **Fig. 6a**). The dissociation between spontaneous recovery of luminance detection (i.e. Humphrey sensitivity) and discrimination performance points towards major differences in the stimuli/tasks necessary to induce different forms of recovery, and in underlying neural mechanisms. It has been postulated that subacute visual defects recover spontaneously as edema and inflammation surrounding the lesion resolve – essentially unmasking networks that were dormant but not destroyed by the stroke (*16–19*). However, the necessity of deliberate training to recover discrimination performance in subacute CB suggests that resolution of edema/inflammation is not sufficient for more complex aspects of visual processing and perception to recover. Such a dissociation may indicate the need to develop more comprehensive clinical tests - beyond perimetry - to assess the extent and depth of visual impairments after occipital strokes and to track patient outcomes in a manner that can better capture the complexities of visual perception.

### Research challenges and future steps towards clinical implementation

The present study was - by necessity of the lack of knowledge of visual properties in subacute CB patients - a non-blinded, laboratory experiment. The discovery at the beginning of our study, as we began testing subacute CB patients, that so many had preserved visual abilities in their blind field was totally unexpected, and required that we alter study goals as the data emerged. As such, our experimental work suffers from limitations associated with the lack of blinding of the investigators/participants and only partial randomization of a subset of the subacute patients (those with global motion deficits) into a training and untrained group. Using the knowledge we have uncovered, future studies can now plan to incorporate a more balanced, unbiased design, as well as a large sample size, to more impartially evaluate and contrast the efficacy of rehabilitation in subacute and chronic stroke patients.

Another limitation of this work, and indeed a major challenge in all rehabilitation research, is the inherent heterogeneity of stroke patients. Stroke lesions – even when restricted to a single vascular territory – are highly variable, as are patient characteristics such as age and comorbidities. We have attempted to control for these factors as much as possible by limiting our study to isolated occipital stroke in patients with otherwise healthy neurologic and ophthalmologic backgrounds. Nonetheless, variability in lesion size, location of visual field deficit, and extent of undamaged, extrastriate visual cortical areas remained (see **Figs. 1 and 2**). As such, though all participants had verifiable V1 damage and training locations were selected and monitored using standard criteria, inconsistencies in training effects may relate to these individual differences.

Additionally, we were unable to identify precisely *when* subacutes lose visual discrimination abilities in the first six months after stroke. It appears that static orientation discrimination is affected before motion discriminations, though we have not measured other form (e.g. shape) discriminations, and do not know when they are lost. As the field of stroke rehabilitation advances, efforts towards clinical implementation will be aided by development of additional metrics and biomarkers to further stratify patients’ recovery potential (*59*). Moreover, while the present study employed a CDDI training program because it was previously identified as highly effective in chronic CBs (*7, 9, 10*), we have yet to determine which training stimulus and paradigm is best suited to treat subacute CBs – ongoing work is investigating this question, as well as the use of adjuvants such as non-invasive brain stimulation, which may further augment the effects of training (*60*). Clinical translation will ultimately depend on all these determinations, as well as the development of services to teach patients how to train properly (especially with accurate fixation), while automatically monitoring their progress and customizing their training locations as needed. Supervised sessions may further extend the clinical utility of training to include patients with cognitive or attentive issues who otherwise would not be able to engage with the program independently.

In summary, subacute CB fields retain a surprisingly large and robust range of visual discrimination abilities. Absent intervention, these are lost by six months post-stroke, possibly a functional consequence of trans-synaptic retrograde degeneration and progressive “*disuse”* of the blind field. Yet, through early training, discrimination abilities can be both retained and further harnessed to recover some of the already-lost visual perception in the blind field. Compared to chronic participants, subacutes trained faster and with greater spatial generalization of learning, both significant advantages for clinical implementation. Fundamentally, our findings challenge the notion that cortically blind fields are a barren sensory domain, and posit that preserved visual abilities indicate rich sensory information processing that temporarily circumvents the permanently-damaged regions of cortex. Thus, after damage to the adult primary visual cortex, judicious, early visual training may be critical to prevent degradation and enhance restoration of preserved perceptual abilities.

## Materials and Methods

### Participants

All participants sustained V1 damage in adulthood, confirmed by neuroimaging, and accompanied by contra-lesional homonymous visual field defects. Additional eligibility criteria included reliable visual fields at recruitment as measured by Humphrey perimetry (see below), and stable, accurate fixation during in-lab, psychophysical, gaze-contingent testing enforced with an eye tracker (see below). Participants were excluded if they had ocular disease (e.g. cataracts, retinal disease, glaucoma), any neurological or cognitive impairment that would interfere with proper training, or hemi-spatial neglect. All participants were best-corrected using glasses or contact lenses. Procedures were conducted in accordance with the Declaration of Helsinki, with written informed consent obtained from each participant, and participation at all times completely voluntary. This study was approved by the Research Subjects Review Board at the University of Rochester.

### Perimetric mapping of visual field defects

Perimetry was conducted using the Humphrey Field Analyzer II-i750 (Zeiss Humphrey Systems, Carl Zeiss Meditec). Both the 24-2 and the 10-2 testing patterns were collected in each eye, using strict quality-control criteria, as previously described (*7*). All tests were performed at the University of Rochester Flaum Eye Institute, by the same ophthalmic technician, with fixation controlled using the system’s eye tracker and gaze/blind spot automated controls, visual acuity corrected to 20/20, a white size III stimulus, and a background luminance of 11.3 cd/m^2^.

Following testing, the four test patterns were interpolated in MATLAB (Mathworks) to create a composite visual field map of each patient, as previously described (*7*). First, luminance detection thresholds were averaged from locations identical in the two eyes. Next, natural-neighbor interpolation with 0.1 deg^2^ resolution was applied between non-overlapping test locations across the four tests, creating composite visual fields of 121 tested locations and 161,398 interpolated data points, subtending an area of 1,616 deg^2^. To determine changes over time, difference maps were generated by subtracting the initial composite visual field map from the subsequent map; change (areas improved or worsened) was defined as visual field locations that differed by at least 6 dB, a conservative standard of change at twice the measurement error of the Humphrey test (Zeiss Humphrey Systems, Carl Zeiss Meditec).

In addition to areas of change across visual field maps, the following measures were collected for each visual field: pattern deviation (Humphrey-derived metric for the deviation from the age-corrected population mean at each HVF test location), perimetric mean deviation (Humphrey-derived metric comparing the overall field of vision to an age-matched normal hill of vision), and total deficit area (defined as regions ≤10 dB of sensitivity, per the standard definition of legal blindness (*61*)).

### Apparatus and eye tracking for in-lab psychophysical measures

Visual discrimination tasks were performed on a MacPro computer with stimuli displayed on an HP CRT monitor (HP 7217A, 48.5×31.5 cm screen size, 1024×640 resolution, 120 Hz frame rate). The monitor’s luminance was calibrated using a ColorCal II automatic calibration system (Cambridge Research Systems) and the resulting gamma-fit linearized lookup table implemented in MATLAB. A viewing distance of 42 cm was ensured using a chin/forehead rest. Eye position was monitored using an Eyelink 1000 eye tracker (SR Research Ltd.) with a sampling frequency of 1000 Hz and accuracy within 0.25°. All tasks and training were conducted using MATLAB (Mathworks) and Psychtoolbox (*62*).

### Visual discrimination testing and training

#### Baseline measurements of visual performance

A battery of two-alternative, forced choice (2AFC) tasks was used to assess visual discrimination performance in-lab, at recruitment. In each task, trials were initiated in a gaze-contingent manner: participants began by fixating on a central spot for 1000ms before a stimulus appeared, accompanied by a tone. If eye movements deviated beyond the 2° x 2° fixation window during the course of stimulus presentation, the trial was aborted and excluded. Participants indicated their responses via a keyboard. Auditory feedback was provided to differentiate correct and incorrect responses. Following each test, participants were asked to describe the appearance of the stimuli in as much detail as possible, or to report if they were unable to sense them at all.

#### Coarse direction discrimination and integration (CDDI) task (Fig. 7c)

After participants initiated a trial through fixation for 1000ms, a random dot stimulus appeared for 500ms in a 5° diameter, circular aperture. The stimulus consisted of black dots moving on a mid-grey background (dot lifetime: 250ms, speed: 10deg/s, density: 3dots/deg^2^). Dots moved globally in a range of directions distributed uniformly around the leftward or rightward vectors (*9, 10*). Participants responded whether the global direction of motion was left- or right-ward. Task difficulty was adjusted using a 3:1 staircase, increasing dot direction range (DR) from 0° to 360° in 40° steps (*9, 10*). Each in-lab session consisted of 100 trials per visual field location. Session performance was fit using a Weibull psychometric function with a threshold criterion of 75% correct to calculate DR thresholds. DR thresholds were then normalized to the maximum range of directions in which dots could move (360°), and expressed as a percentage using the following formula: CDDI threshold (%) = [360° - DR threshold]/360° x 100. For ease of analysis, when participants performed at chance (50% correct for a given session), the CDDI threshold was set to 100%.

#### Fine direction discrimination (FDD) task (Fig. 7d)

Random dot stimuli were presented in which black dots (parameters as in CDDI task) moved on a mid-grey background within a 5° circular aperture, for 500ms. Dots moved almost uniformly (2° DR) leftwards or rightwards, angled upward or downward relative to the stimulus horizontal meridian. Participants indicated if the motion direction was up or down. Task difficulty was adjusted using a 3:1 staircase, which decreased angle of motion from the horizontal meridian from 90° (easiest) to 1° in log steps. Each test session consisted of 100 trials at a given location. Session performance was fit using a Weibull function with a threshold criterion of 75% correct to calculate fine direction difference (FDD) thresholds. At chance performance (50% correct), the FDD threshold was set to 90°.

#### Contrast sensitivity functions (CSFs) for orientation and direction discrimination

CSFs were measured using the quick CSF (qCSF) (*63*), a widely used Bayesian method to measure the entire CSF across multiple spatial frequencies (0.1-10 cycles/deg) in just 100 trials, with a test-retest reliability of 0.94 in clinical populations (*64*). In this method, the shape of the CSF is expressed as a truncated log-parabola, defined by four parameters: peak sensitivity, peak spatial frequency, bandwidth, and low-frequency truncation level. In the motion qCSF (**Fig. 7e**), participants performed a 2AFC left *versus* right, motion direction discrimination task of a vertically-oriented Gabor (5° diameter, sigma 1°, 250 ms on/off ramps). In our static qCSF (**Fig. 7f**), participants had to discriminate horizontal from vertical orientation of a non-flickering Gabor patch (5° diameter, sigma 1°, 250 ms on/off ramps). Velocity varied as a function of spatial frequency to ensure a temporal frequency of 10 Hz (*63*).

**Fig. 7.**
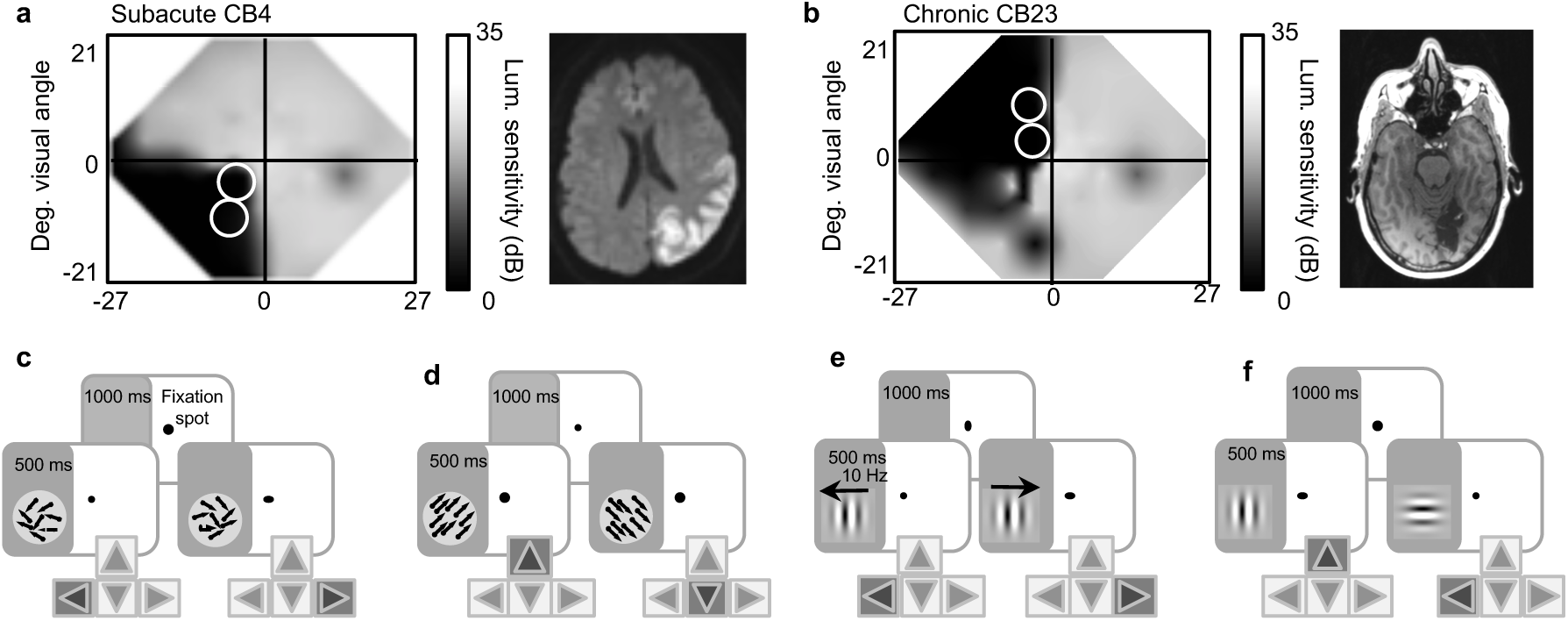
Measuring and retraining vision in subacute and chronic stroke. **a**, Interpolated Humphrey Visual Field (HVF) and diffusion-weighted magnetic resonance image (MRI) of the brain of subacute participant CB4 (2.5 months post-stroke). White circles on the HVF denote the location/size of testing/training stimuli in the blind field. Brain image is veridically oriented (right hemisphere is on the right of image). **b**, Interpolated HVF, testing/training locations and T1-weighed MRI of chronic participant CB23 (29 months post-stroke). Note clear difference in the presentation of the brain damage (including shrinkage of the occipital grey and white matter, ventricular enlargement) compared to the subacute participant in **a**. **c-f**, Trial sequences for tasks measuring coarse direction discrimination and integration (CDDI) (**c**), fine direction discrimination (FDD) (**d**), and contrast sensitivity for direction (**e**) and static orientation discrimination (**f**).

#### Visual training in cortically blind fields

after Humphrey perimetry, each subject underwent psychophysical mapping of the blind field border as previously described (*9, 10*). Training locations were selected as sites where performance first declined to chance (50-60% correct, CDDI threshold=100%) during mapping; two initial training locations were identified in each subject, including those subsequently assigned to the untrained group.

Following selection of putative training locations, 8 subacute and 14 chronic CB participants were assigned to train on the CDDI task. Participants used their personal computers and displays to perform daily training at home. They were supplied with chin/forehead rests and instructed to position them such that their eyes were 42 cm away from their displays during training. They performed 300 trials per training location per day, at least five days per week, and emailed their auto-generated data log files back to the laboratory for analysis on a weekly basis. Session thresholds were calculated by fitting a Weibull function with a threshold criterion of 75% correct performance. DR thresholds were converted to CDDI thresholds as described above. Once performance reached levels comparable to equivalent, intact field locations, training moved 1° deeper into the blind field along the x-axis (Cartesian coordinate space). Although home-training was performed without an eye tracker, patients were instructed to fixate whenever a fixation spot was present and warned that inadequate fixation would prevent recovery. Once subacute participants reached the chronic period, they were brought back to Rochester and performance at all home-trained locations was verified in-lab with on-line fixation control using the Eyelink 1000 eye tracker (SR Research). Chronic participants’ performance post-training was similarly verified in lab with eye tracking.

### Statistical Analyses

To evaluate differences in threshold performance for the CDDI and FDD tasks, when three or more groups were compared, inter-group differences were tested with a one- or two-way analyses of variance (ANOVA) followed by Tukey’s post-hoc tests, if appropriate. When only two groups were compared, a two-tailed Student’s t-test was performed. Repeated measures statistics were used whenever appropriate. A probability of type I error of *p*<0.05 was considered statistically significant.

Due to the adaptive nature of the qCSF procedure, a bootstrap method (*65*) was used to determine statistically significant changes to the CSF across experimental conditions and groups. Specifically, the qCSF procedure generates an updated prior space for each of four CSF parameters on each successive trial. To generate a bootstrapped distribution of CSF parameters (and, by extension, contrast sensitivities at each spatial frequency) we performed the following procedure: For each experimental run, we generated 2,000 CSFs by running the qCSF procedure 2,000 times using 100 trials randomly sampled with replacement from the data collected at that location. To compute p-values for comparisons of CSFs, we compiled difference distributions for the comparisons in questions for each model parameter and contrast sensitivities at each tested spatial frequency, with p-values determined by the proportion of samples that “crossed” zero. To estimate a floor for a case of no visual sensitivity, we simulated 10,000 qCFS using random responses. In this simulation, the 97.5 percentile peak CSF sensitivity was 2.55. For actual patient data, any sample with CSF with peak sensitivity of less than 2.55 was considered at chance level performance, and thus set to zero. This ensured that we used a conservative criterion for determining whether patients exhibited contrast sensitivity in their blind filed. Then, using bootstrap samples, we obtained 95% confidence intervals for CSF parameters and contrast sensitivities for individual spatial frequencies as well as associated p-values.

## Acknowledgments

The authors wish to thank Terrance Schaefer, who performed Humphrey visual field tests presented here. We also thank Drs. Shobha Boghani, Steven Feldon, Alexander Hartmann, Ronald Plotnick, Zoe Williams (Flaum Eye Institute); Ania Busza and Bogachan Sahin (Department of Neurology) for patient referrals. Some of the data in chronic patients analyzed here were collected by Anasuya Das and Matthew Cavanaugh as part of two prior studies.

## Funding

The present study was funded by NIH (EY027314 and EY021209, T32 EY007125 to Center for Visual Science, T32 GM007356 to the Medical Scientist Training Program, TL1 TR002000 to the Translational Biomedical Sciences Program, a pilot grant from University of Rochester CTSA award UL1 TR002001 to ELS), and by an unrestricted grant from the Research to Prevent Blindness (RPB) Foundation to the Flaum Eye Institute.

## Author contributions

ELS and KRH designed the study; ELS collected the data; ELS, KRH, MM, and DT analyzed the data; ELS wrote the manuscript, and all authors commented on it.

## Competing interests

KRH is co-inventor on US Patent No. 7,549,743 and has founder’s equity in Envision Solutions LLC, which has licensed this patent from the University of Rochester. The University of Rochester also possesses equity in Envision Solutions LLC. The remaining authors have no competing interests.

## Data and materials availability

All de-identified data are available from the authors upon request.

